# Siglec-5 is an inhibitory immune checkpoint molecule for human T cells

**DOI:** 10.1101/2022.01.04.474948

**Authors:** Aleksandra Vuchkovska, David G. Glanville, Gina M. Scurti, Paula White, Michael I. Nishimura, Andrew T. Ulijasz, Makio Iwashima

## Abstract

Sialic acid-binding immunoglobulin-type lectins (Siglecs) are a family of immunoglobulin-type lectins that mediate protein-carbohydrate interactions via sialic acids attached to glycoproteins or glycolipids. Most of the CD33-related Siglecs (CD33rSiglecs), a major subfamily of rapidly evolving Siglecs, contain a cytoplasmic signaling domain consisting of the immunoreceptor tyrosine-based inhibitory motif (ITIM) and immunoreceptor tyrosine-based switch motif (ITSM) and mediate suppressive signals for lymphoid and myeloid cells. While most CD33rSiglecs are expressed by innate immune cells, such as monocytes and neutrophils, to date, the expression of Siglecs in human T cells has not been well appreciated. In this study, we found that Siglec-5, a member of the CD33rSiglecs, is expressed by most activated T cells upon antigen receptor stimulation. Functionally, Siglec-5 suppresses T cell activation. In support of these findings, we found that Siglec-5 overexpression abrogates antigen receptor induced activation of Nuclear factor of activated T cells (NFAT) and Activator protein 1 (AP-1). Furthermore, we show that GBS β-protein, a known bacterial ligand of Siglec-5, reduces the production of cytokines and cytolytic molecules by activated primary T cells in a Siglec-5 dependent manner. Our data also show that some cancer cell lines express a putative Siglec-5 ligand(s), and that the presence of soluble Siglec-5 enhances tumor-cell specific T cell activation, suggesting that some tumor cells inhibit T cell activation via Siglec-5. Together, our data demonstrate that Siglec-5 is a previously unrecognized inhibitory T cell immune checkpoint molecule and suggests that blockade of Siglec-5 could serve as a new strategy to enhance anti-tumor T cell functions.

**One Sentence Summary:** Siglec-5 is a novel checkpoint receptor expressed by activated human T cells that antagonizes TCR mediated activation and suppresses T cell effector functions such as cytokine production.

## INTRODUCTION

To prevent inappropriate immune activation and subsequent immune injuries, the immune system has evolved strategies to help discriminate self from non-self, or healthy from altered and infected. These strategies rely on the activation of different receptors that recognize specific molecular patterns to differentiate between self and healthy cells (*1–3*). Sialic-acid-binding immunoglobulin-like lectins (Siglecs), a family of C-type lectins that binds sialylated glycans, are a group of receptors that are mainly expressed by innate immune cells and have immunomodulatory functions (*4*). Sialic acids are derivatives of the sugar neuraminic acid and are attached on the terminal position of glycoproteins or glycolipids. When sialic acids engage the Siglec family of receptors, they modulate the activation and effector functions of Siglec expressing immune cells (*5*). Most Siglecs of the rapidly evolving CD33 related subfamily (CD33rSiglec) have immune suppressive functions which they mediate through the conserved immunoreceptor tyrosine-based inhibitory motifs (ITIMs) or immunoreceptor tyrosine-based switch motifs (ITSMs)(*4*). The arrangement of the ITIM and ITSM of most inhibitory Siglecs mirrors the arrangement in the well-known inhibitory checkpoint receptor PD-1 (*6*). However, little to no expression of CD33rSiglecs has been reported for T cells, with T cell CD33rSiglec expression only found in the context of some pathological conditions. For example, HIV infected patients have a greater proportion of circulating CD4^+^ T cells expressing Siglec-5 and Siglec-9 as compared to uninfected healthy donors. Expression of these inhibitory Siglecs correlates with resistance to excessive immune activation and subsequent HIV-induced cell death (*7*). Similarly, tumor infiltrating lymphocytes (TILs) from various cancers, including melanoma and non-small cell lung cancer (NSCLC), express several inhibitory CD33rSiglecs, such as Siglec-3, Siglec-5, Siglec-7, Siglec-10, and Siglec-9. Importantly, Siglec-9 expression in TILs negatively correlates with the prognosis and survival of NSCLC patients (*8, 9*).

Here we demonstrate that human activated T cells express Siglec-5. Previous work by others showed that Siglec-5 suppresses the immune responses of innate immune cells such as monocytes and neutrophils against pathogens (*10, 11*). For example, some strains of Group B Streptococcus (GBS) engage Siglec-5 via the surface *β*-protein, an action that reduces bacterial clearance by suppressing bacterial killing and phagocytosis (*12, 13*). In adults, GBS is a common commensal bacterium in the gastrointestinal and vaginal tract; however, in newborns, GBS is the leading cause of life-threatening sepsis and meningitis (*14*). As seen with innate immune cells, our data show that Siglec-5 has a previously unrecognized and novel inhibitory function during T cell activation. In this work we demonstrate this novelty by showing that overexpression of Siglec-5 inhibits the activity of T cell receptor (TCR)-induced transcription factors, such as NFAT and AP-1. Furthermore, engagement of Siglec-5 with the specific GBS *β*-protein ligand reduces primary T cell activation and leads to decreased production of cytokines and activation-induced effector molecules. The data also show that some cancer cell lines express putative ligands that bind Siglec-5. Importantly, T cell responses against the putative ligand positive cancer cell lines are enhanced in the presence of soluble Siglec-5. Together, our data show that Siglec-5 is a novel and previously unidentified inhibitory T cell immune checkpoint molecule, which could pave the way for novel immune therapies.

## RESULTS

### CD33rSiglec expression by human T cells

Innate immune cells such as monocytes and neutrophils express multiple CD33rSiglecs (*4*). In contrast, little to no expression is reported in T cells (*15*), with the exception of Siglec-7 and -9, which are expressed by small populations of CD8+ T cells and mediate direct inhibition of TCR signaling (*16*). To obtain a global perspective on Siglec expression by T cells, we assessed the expression of the CD33rSiglec family members (Siglec-3, -5, -7, -8, -9 and -10) in both resting and stimulated cells. As a positive control for the staining, we also included human monocytes from adult peripheral blood mononuclear cells (PBMCs) (Fig.S1). Among the Siglecs we tested, Siglec-5 showed a robust activation-associated pattern of expression by both CD4 and CD8 T cells (Fig.1A and 1B). In contrast, we observed minimal levels of Siglec-6, -8, -9, and -10 expression and a low level of Siglec-3 (both CD4 and CD8) and Siglec-7 (CD8) expression (Fig.1A). Siglec-3 and Siglec-7 expression in the T cells was significantly lower compared to Siglec-5 and did not demonstrate activation-associated pattern of expression. As a result of these observations, we therefore focused on the expression and function of Siglec-5 in human T cells.

**Fig. 1.**
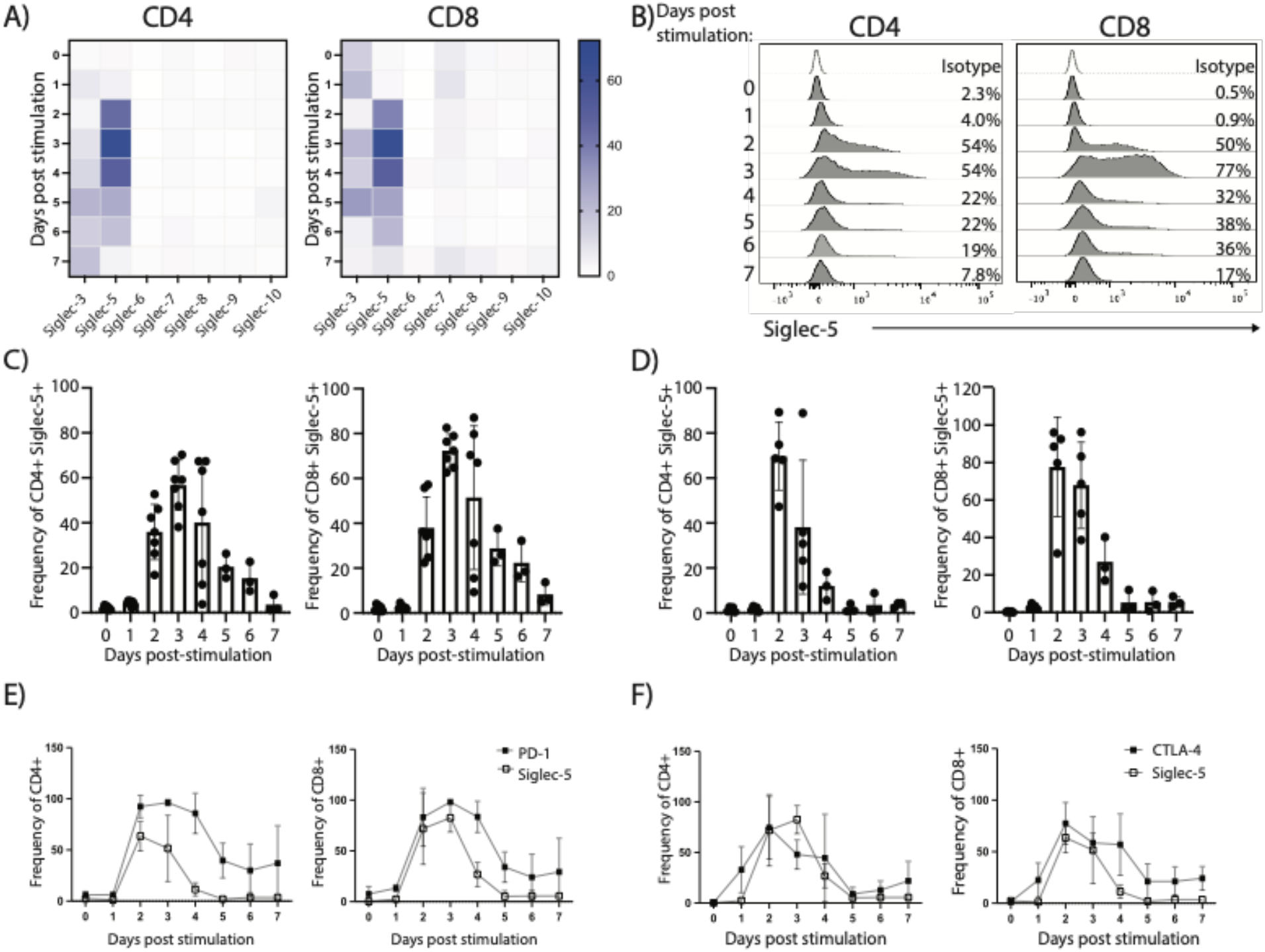
CD33rSiglec expression in resting and activated T cells. Adult peripheral blood or cord blood mononuclear cells were subjected to *in vitro* activation using soluble anti-CD3 and IL-2 for up to 7 days. Cells were split every 2-3 days. (**A**) Expression of Siglecs-3, -5, -6, -7, -8, -9, -10 was evaluated in the CD4 and CD8 T cell from multiple adult donors. (**B**) Representative plots and (**C**) summary for multiple adult donors (each dot represents an individual donor) of Siglec-5 expression within CD4 and CD8 T cells. (**D**) Summary for multiple cord blood donors of Siglec-5 expression within CD4 and CD8 T cells. (**E**) Summary for multiple cord blood donors (n=3) for expression of Siglec-5 and PD-1 and (**F**) Siglec-5 and CTLA-4 within CD4 and CD8 T cells.

Cell surface expression of Siglec-5 was detected at 48hrs post activation in both adult and cord blood CD4 and CD8 T cells (Fig.1C and 1D). In adult T cells, Siglec-5 expression peaked at 72hrs post-activation, upon which it begun to gradually decrease and was lost by day 7 post-activation (Fig.1B and 1C). Cord blood T cells also demonstrated Siglec-5 expression after stimulation. The surface expression of Siglec-5 by cord blood T cells reached peak expression earlier than in adult T cells, at 48hrs post stimulation (Fig.1D). Siglec-5 expression kinetics overlapped with the co-inhibitory checkpoint receptor PD-1 (Fig.1E and Fig.S2A), while an additional co-inhibitory checkpoint receptor, CTLA-4, preceded Siglec-5 and its expression started one day post-stimulation (Fig.1F and Fig.S2B). Similarly, Siglec-5’s expression kinetics closely overlapped with OX-40 expression (Fig.S3B), but lagged behind ICOS expression, which peaks at day 1 post-stimulation (Fig.S3A). 4-1BB expression is much lower in CD4 T cells compared to Siglec-5, and CD8 T cells express 4-1BB earlier than Siglec-5 (Fig.S3C).

### Post-translational regulation of surface expression of Siglec-5

We next tested if Siglec-5 expression is regulated via transcriptional and/or post-translational mechanisms. We performed quantitative RT-PCR to determine the amount of *siglec-5* mRNA in resting and stimulated adult T cells. In contrast to the surface expression, peak mRNA expression of *siglec-5* was observed within 24 hours after stimulation and rapidly decreased after that (Fig.2A). The time gap between mRNA and surface Siglec-5 expression suggested that there is a post-transcriptional process that controled the surface expression of Siglec-5. Similar regulation has been observed for CTLA4, which surface expression is tightly controlled by membrane endo/exocytosis (*17*). To test how Siglec-5 surface expression is controlled, we determined Siglec-5 protein levels via Western blot. Surface expression of Siglec-5 was not detectable on the surface until 48hrs post-stimulation (Fig.1B and 1C), however Siglec-5 protein was clearly present in activated T cells as early as 24hrs post-stimulation (Fig.2B). These data suggest that there is a post-translational mechanism that controls the surface expression of Siglec-5, possibly by membrane trafficking and/or endo/exocytosis of the protein to the cell surface.

**Fig. 2.**
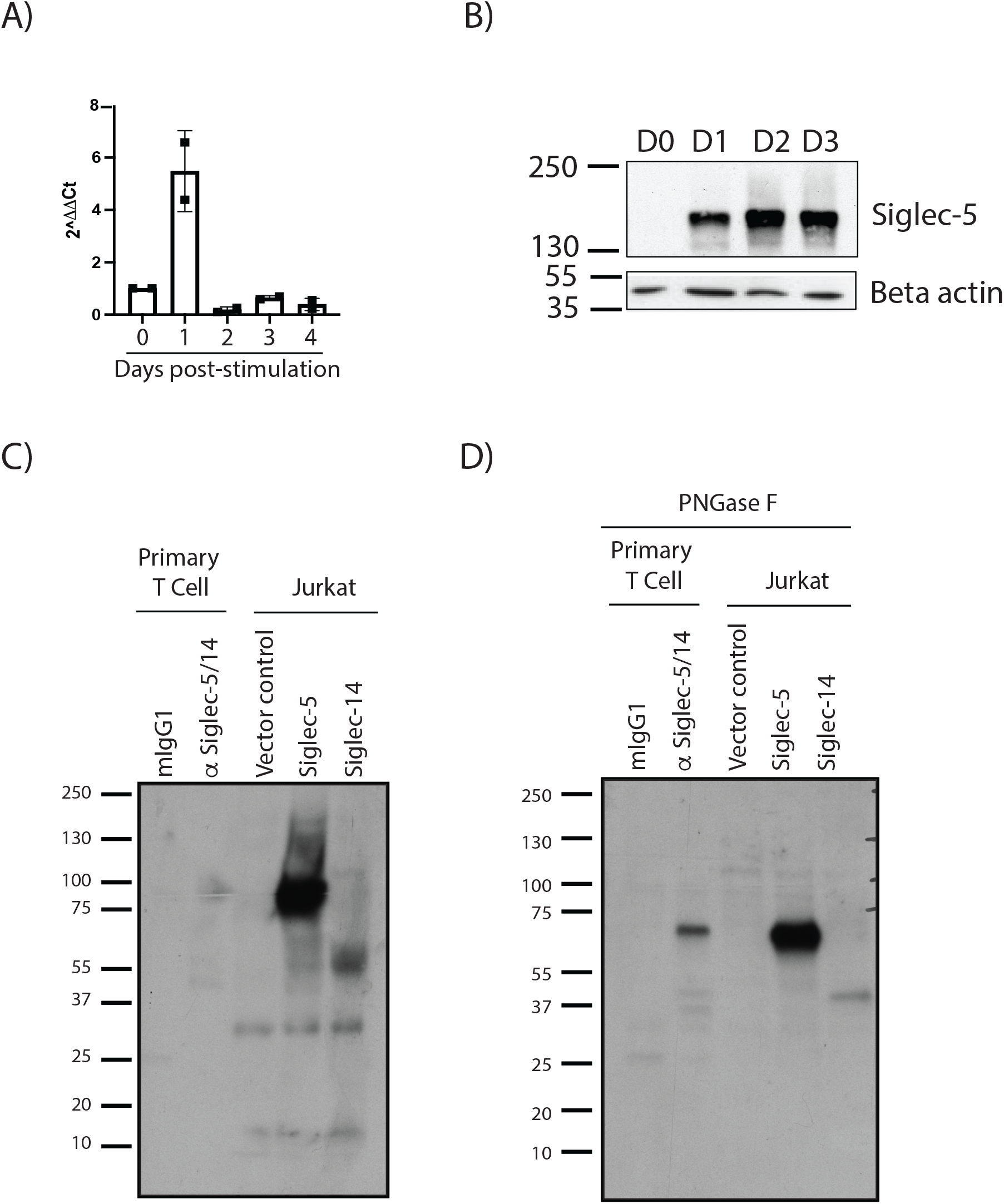
Verification of Siglec-5 expression in T cells. (**A**) CD3+ T cells were enriched from adult PBMCs and cultured with plate bound anti-CD3 and anti-CD28 stimulation and IL-2, for up to 4 days. mRNA was prepared at each time point and Siglec-5 mRNA was detected using qPCR. (**B**) Representative blot (n=3) of CD3+ T cells enriched from adult PBMCs cultured with plate bound anti-CD3 and anti-CD28 stimulation and IL-2, for up to 3 days. At each time point, cell lysates were prepared using denaturing non-reducing conditions. (**C**) and (**D**) Cord blood T cells were stimulated with soluble anti-CD3 and IL-2 for 2 days. Cells were lysed with mild detergent (0.5% NP-40) and proteins were immunoprecipitated (IP) using anti-Siglec-5/14 mAb (clone 1A5) or mIgG1 isotype CNBr conjugated beads. As controls, Jurkat T cells were transfected with Siglec-5, Siglec-14 or empty vector and whole cell lysates were prepared. Where indicated, IP-ed proteins were treated with PNGase F to remove N-linked glycosylation. All samples were run on SDS-PAGE and transferred to PVDF membranes. Membranes were blotted with anti-Siglec5/14 polyclonal antibody.

There is a known Siglec family member, Siglec-14, that shares 74% protein sequence homology with Siglec-5 within the ectodomain (*18*). Most commercially available antibodies cross-react and recognize both Siglec-5 and Siglec-14. Thus, it is important to confirm that the anti-Siglec-5 antibody (clone 1A5) used for flow cytometric analysis of the surface protein (Fig.1) detects Siglec-5 and not Siglec-14 following T cell activation. As shown in Fig.2B, using samples from activated human T cells and prepared under non-reducing conditions we detect a major protein band at molecular weight (MW) 150kD. This band corresponds to the predicted homodimer formed by Siglec-5 (*10*). To further confirm that the protein we detected is Siglec-5 and not Siglec-14, we immunoprecipitated proteins from activated primary T cells using anti-Siglec-5/-14 antibody (clone 1A5). As a control, we overexpressed Siglec-5 or Siglec-14 cDNA in Jurkat T cells. The predicted size of the Siglec-5 monomer is 68kDa, while that of Siglec-14 is 42kD. The bands we observed from the immunoprecipitated samples (using polyclonal anti-Siglec-5/-14 antibody) corresponded to our control Siglec-5 overexpression in Jurkat T cells.

However, both the control and the immunoprecipitated bands ranged around 85kD (Fig.2C), a higher MW than the predicted 68kDa for Siglec-5. Because Siglec-5 has at least eight described N-based glycosylation sites (*19*), we hypothesized that the higher-than-expected MW is due to the glycosylation. Indeed, when we treated the samples with PNGase F to remove N-linked glycosylation, we observed that the bands from the control Siglec-5 overexpressing Jurkat T cells and immunoprecipitated proteins from primary T cells corresponded to the predicted 68kDa band for Siglec-5 (Fig.2D). We also observed a lower MW band in the immunoprecipitated samples from primary T cells, but these bands did not migrate at the same predicted molecular weight as the band from control Siglec-14 overexpressing Jurkat T cells. These data suggest that the bands are likely from other proteins that have cross reacted with the blotting antibodies.

### Function of Siglec-5 in T cells

Studies evaluating Siglec-5 expression in innate immune cells such as monocytes and neutrophils show that Siglec-5 is an inhibitory receptor that mediates its function through the recruitment of Shp1 and Shp2 phosphatases to the ITIM and ITSM present in the cytoplasmic region of Siglec-5 (*20*). Among other inhibitory Siglecs, Siglec-5 has the highest level of similarity with PD-1 within the signaling domains (ITIM and ITSM) and the spacer (Fig.3A). Based on its similarities with PD-1, as well as its previously described function in innate immune cells, we hypothesized that Siglec-5 could be a negative regulator of T cell activation. To test this, we assessed if overexpression of Siglec-5 could reduce TCR-induced transcription factor activation by co-transfecting a Siglec-5 expression vector with NFAT and AP-1 luciferase reporter vectors and measuring the luciferase activity with and without TCR stimulation. Indeed, overexpression of Siglec-5 significantly reduced TCR-induced NFAT and AP-1 activity (Fig.3B), which is consistent with our hypothesis that Siglec-5 is a novel inhibitory molecule in T cells. Since ITIM and ITSM are both known to mediate the suppressive functions of PD-1 (21), we hypothesized that Siglec-5 also requires these motifs to suppress NFAT and AP-1. To test this, we generated a truncated Siglec-5 (ΔSiglec-5) that lacks both ITIM and ITSM. If these motifs are required for suppression, we expected to see restoration of NFAT activity in Jurkat T cells expressing ΔSiglec-5. Indeed, we observed a partial restoration in NFAT activity by cells expressing ΔSiglec-5 (Fig.3C). Interestingly, expression of ΔSiglec-5 caused an increase in NFAT activity even in unstimulated cells (Fig.3C), suggestive of an unregulated constitutive function.

**Fig. 3.**
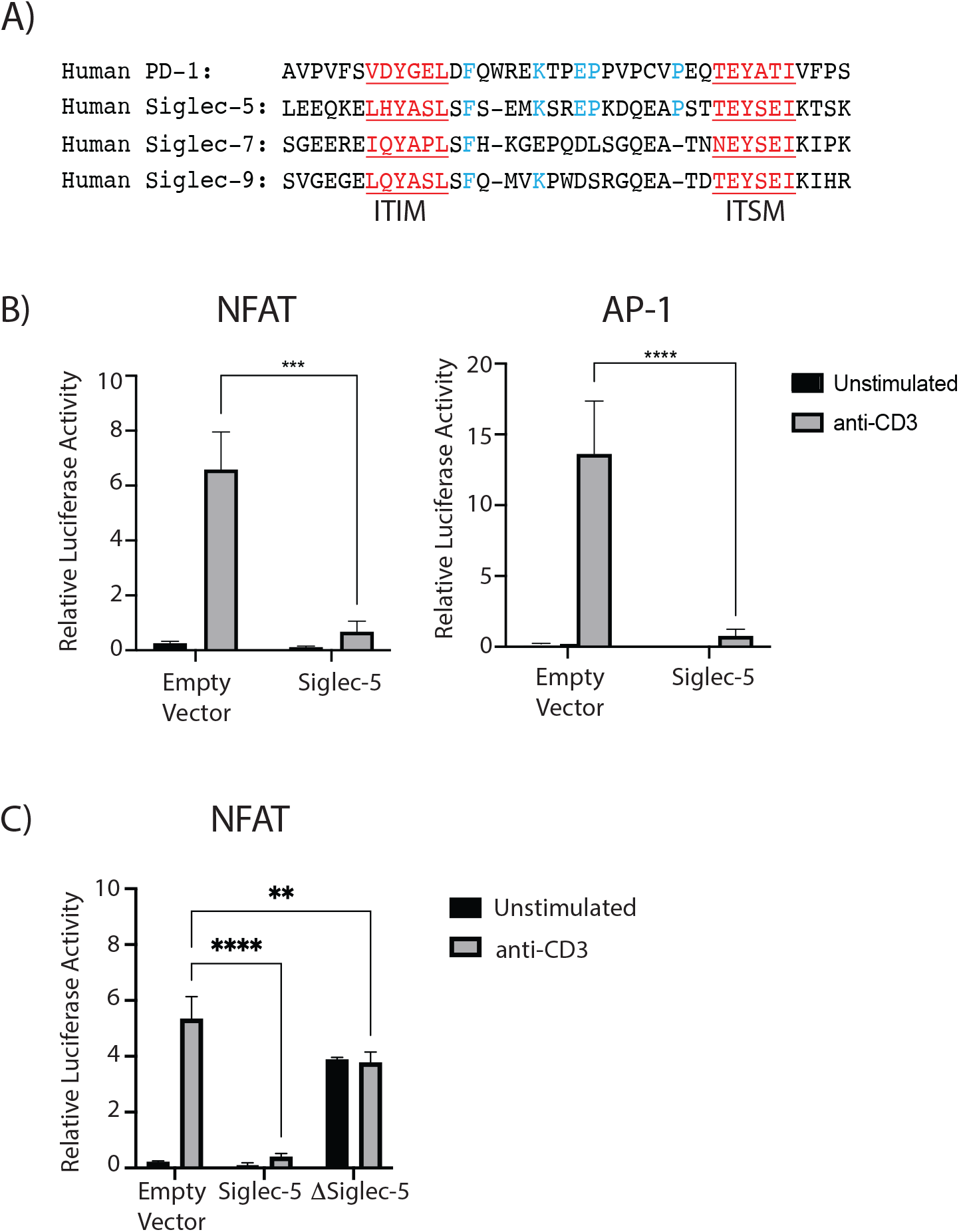
Function of Siglec-5 in T cells. (**A**) Sequence alignment of the ITIM and ITSM domains of PD-1 and Siglec-5, -7 and -9. Shared protein motifs are highlighted in red, and conserved amino acids are highlighted in blue. (**B**) and (**C**) Jurkat T cells were transiently transfected with NFAT or AP-1 luciferase reporter vectors, along with A) full length Siglec-5 or B) truncated Siglec-5 (lacking ITIM and ITSM domains). 4hrs post anti-CD3 stimulation, NFAT and AP-1 luciferase activity was measured. Relative luciferase activity was calculated based on Renilla luciferase activity. Representative figures of three independent experiments. Statistical analysis: 2wayANOVA, Tukey’s multiple comparison test, *** p<0.001, **** p<0.0001.

### Role of Siglec-5 in Group B Streptococcus β-protein mediated suppression of T cell activation

Pathogens have evolved to evade the immune responses by utilizing immunomodulatory mechanisms that normally serve to keep immune cells from over activation (*22*). Some strains of Group B Streptococcus (GBS), a major pathogen of newborns (*14*), express a surface protein called *β*-protein, which is known to suppress the immune response via binding to Siglec-5 in innate immune cells (*23*). The Siglec-5 binding region of the GBS *β*-protein resides within the N-terminal 152aa and is termed the B6N domain (*12*). *β*-protein/Siglec-5 interactions result in inhibition of phagocytosis and suppression of proinflammatory cytokine production, such as IL-8, by neutrophils and monocytes in response to GBS (*12,13,23*). To address the function of Siglec-5 during T cell activation, we tested whether engaging Siglec-5 with the B6N region of the *β*-protein would inhibit primary T cells activation. We generated a B6N fusion protein with superfolder (sf) Green Fluorescent Protein (GFP) (termed B6N::sfGFP) (Fig.S4A). The binding capability of Siglec-5 to B6N::sfGFP, but not control sfGFP, was confirmed using an indirect enzyme-linked immunoassay (ELISA) (Fig.S4B). To test if Siglec-5 suppressed T cell activation, we stimulated human CD4 or CD8 T cells in the presence of the Siglec-5 ligand B6N::sfGFP. As shown in Fig.1, Siglec-5 expression is elevated at 3 days post-stimulation. Thus, we stimulated naïve T cells for 3 days using plate bound anti-CD3 and anti-CD28 antibodies. Three days after stimulation, we harvested the cells and restimulated them with plate bound anti-CD3 and anti-CD28 in the presence of B6N::sfGFP, or the control sfGFP, for an additional 3 days. After the secondary stimulation, we assessed cytokine production and effector molecule expression by T cells. Stimulation of CD4 T cells in the presence of B6N::sfGFP reduced production of several proinflammatory cytokines, such as IFN-*γ* and IL-22 (we did not observe IL-17A production under these conditions) (Fig.4A), while it increased production of Th2 type cytokines such as IL-4, IL-5, IL-13 (Fig. 4B). Similarly, B6N::sfGFP reduced production of IFN-*γ* by CD8 T cells (Fig.4A). Furthermore, B6N::sfGFP also reduced the expression of Granzyme B by activated CD4 T cells (Fig.4C). To test whether B6N::sfGFP changed T cell cytokine production in a Siglec-5 dependent manner, we pre-incubated B6N::sfGFP with soluble Siglec-5 Fc chimeric protein before coating it onto plates to reduce the availability of B6N::sfGFP that can bind the endogenous Siglec-5 molecule on T cells. Indeed, when pre-treated with the Siglec-5 Fc protein, B6N::sfGFP did not suppress the activation of T cells, as measured by the expression Granzyme B and IFN-*γ* (Fig.4D), and demonstrating specificity of B6N::sfGFP for Siglec-5.

**Fig. 4:**
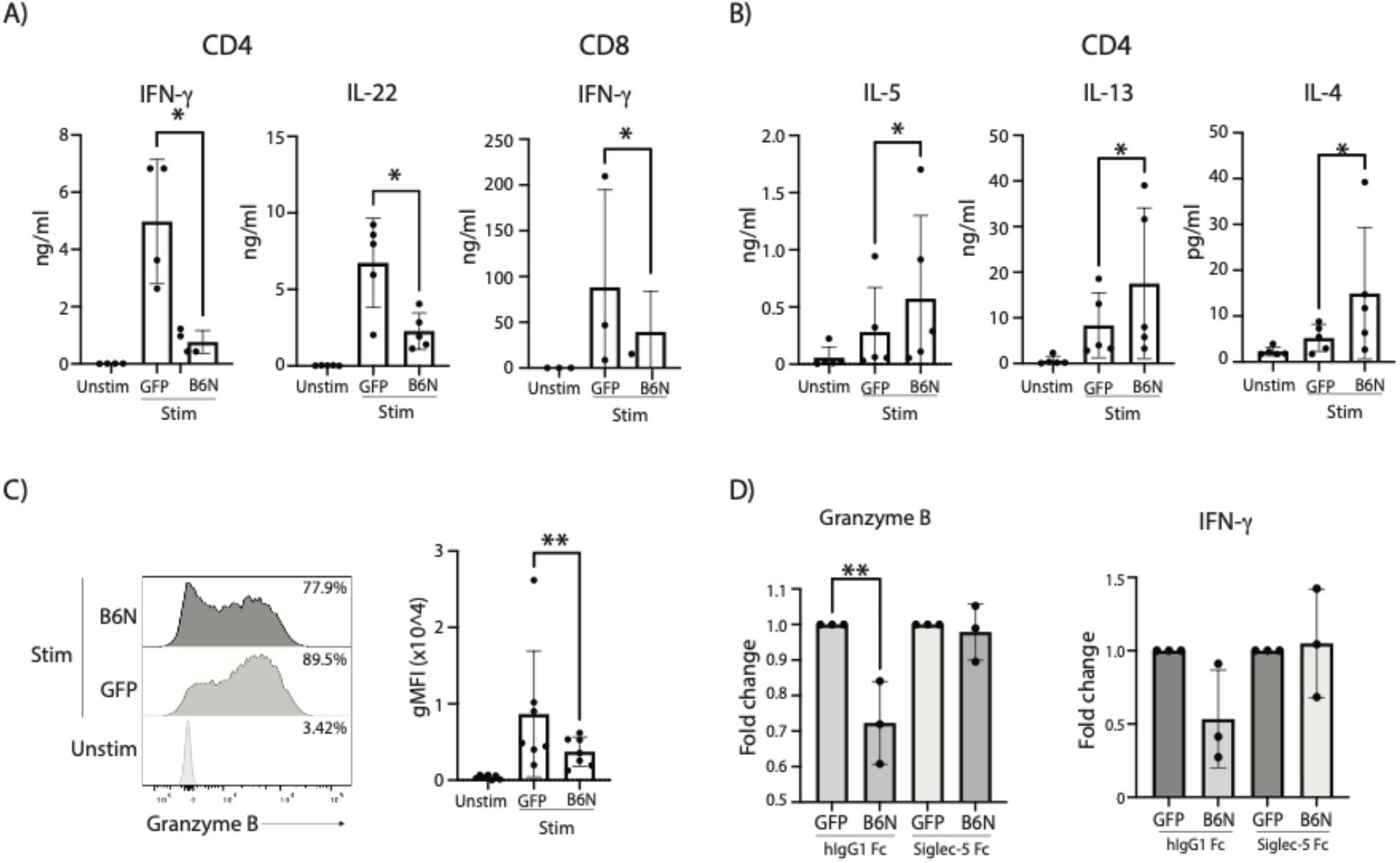
Engagement of Siglec-5 with β-protein and its effect on T cell effector functions. Naïve CD4 or CD8 T cells were cultured with plate bound anti-CD3 and anti-CD28 stimulation in the presence of IL-2 for 3 days. Cells were than harvested and re-stimulated with plate bound anti-CD3, anti-CD28 and B6N::sfGFP or sfGFP. Supernatants and cells from the 3 days cultures were collected and used for (**A**) and (**B**) cytokine bead array analysis and (**C**) measurement of Granzyme B production. Statistical analysis: Ratio paired two-tailed t test; * p<0.05, ** p<0.01. (**D**) Day 3 stimulated CD4 T cells were re-stimulated with pate bound anti-CD3, anti-CD28 and B6N::sfGFP or sfGFP pre-incubated with hIgG1 Fc or Siglec-5 Fc at equimolar ratios. At day 3 post-re-stimulation, cytokine production and Granzyme B expression were measured. For each condition, treatments were normalized to unstimulated values first. Fold change was than calculated as the ration of cells stimulated in the presence of B6N::sfGFP vs sfGFP. Statistical analysis: one-way ANOVA, Tukey’s multiple comparisons test, ** p<0.01. Each dot represents an individual donor.

### Role of Siglec-5 engagement during the T cell specific anti-tumor response

Siglec-5 expression by tumor infiltrating lymphocytes has been reported (*9*). Considering the immunosuppressive functions that inhibitory checkpoint receptors play in anti-tumor immunity, we asked whether Siglec-5 is also involved in the process by which cancers suppress T cell responses. We first investigated if Siglec-5 could interact with human cancer-derived cell lines. Our data showed that this was the case, whereby Siglec-5 interacted with multiple cancer cell lines, including MEL624, Raji and SUN-475, among others, suggesting that these cell lines express putative ligands for Siglec-5 (Fig.S5). To test if the putative ligands expressed by cancer lines could suppress the T cell response, we used 1383i TCR+ T cells. 1383i TCR+ T cells are engineered human T cells transduced with a TCR specific for the melanoma antigen tyrosinase (*24*) and used in clinical trials as T cell therapy for metastatic melanoma (ClinicalTrials.gov ID NCT01586403). First, we tested whether 1383i TCR+ T cells stimulated in an antigen specific manner were able to upregulate Siglec-5 expression. When 1383i TCR+ T cells were stimulated with antigen presenting cells (T2 cells) pulsed with tyrosinase peptide, we observed a robust expression of Siglec-5 (Fig.S6). We hypothesized that a soluble form of Siglec-5 (Siglec-5 Fc) would bind the putative Siglec-5 ligands expressed by MEL624 and interfere with the engagement of the endogenous Siglec-5 molecule expressed by activated cancer specific T cells. To test our hypothesis, we stimulated 1383i TCR+ T cells with irradiated T2 cells pulsed with tyrosinase. Two days post primary stimulation, when Siglec-5 expression is high, we re-stimulated the 1383i TCR+ T cells with the MEL624 cancer line in the presence or absence of Siglec-5 Fc protein. Stimulation with MEL624 pretreated with Siglec-5 Fc led to an increased frequency of IL-2, IFN-*γ*, and TNF-*α*, producing 1383i TCR+ T cells (Fig. 5A). Moreover, CD107a, the surface antigen associated with the release of cytotoxic granules, was also significantly increased by 1383i TCR+ T cells stimulated by MEL624 pre-treated with Siglec-5 Fc (Fig. 5B). In this assay, CD8 T cells re-stimulated with MEL624 did not produce any amounts of cytokines. Presumably due to the anergic state caused by the repeated stimuli. Together, these data showed that engagement of Siglec-5 by putative ligands expressed on the target cells suppressed CD4 T cell activation and further supported our hypothesis that Siglec-5 functions as a novel inhibitory checkpoint molecule for activated human T cells.

**Fig. 5.**
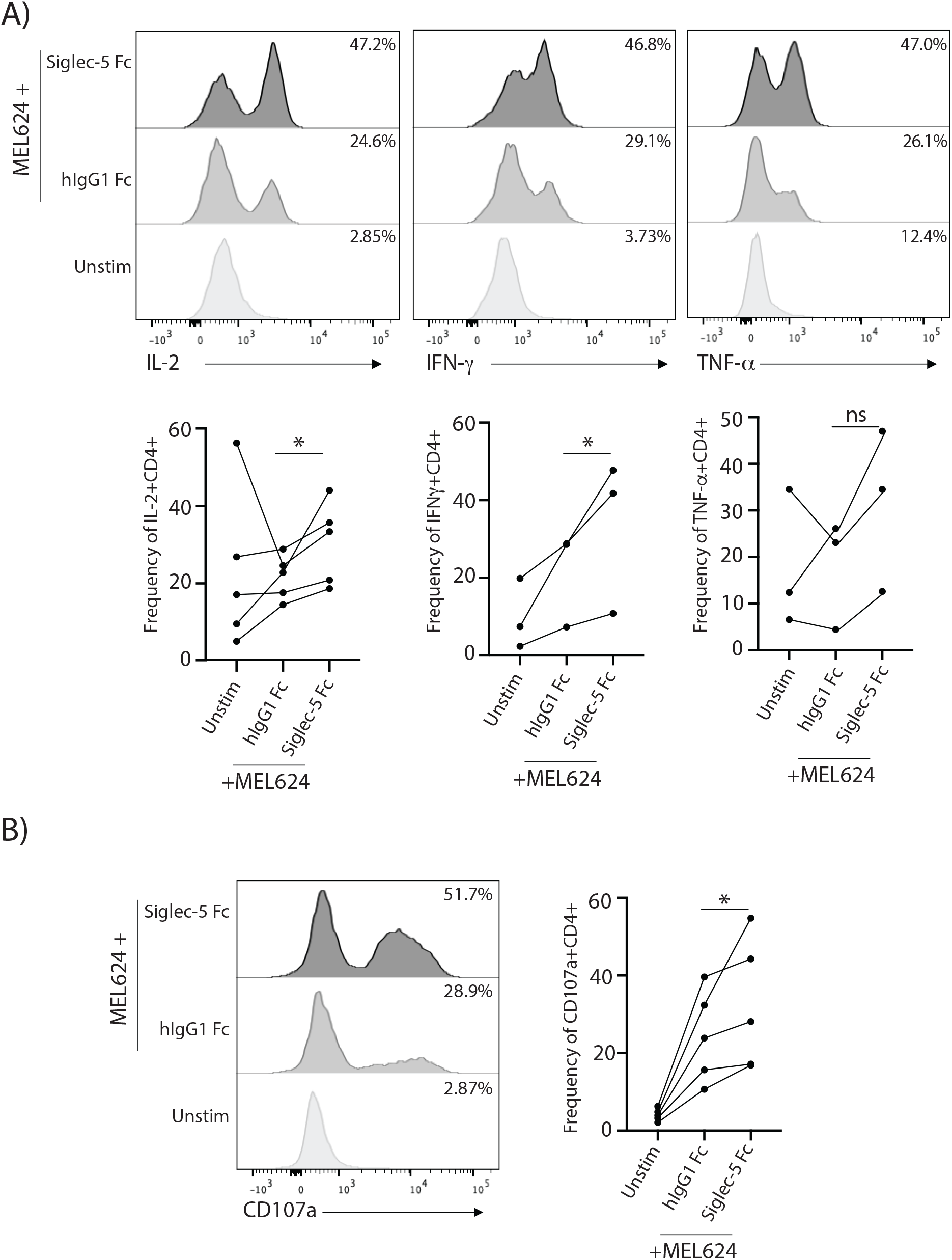
Disrupting the Siglec-5 signaling axis using soluble Siglec-5 Fc. 1383i T cells were stimulated with T2 pulsed with tyrosinase peptide. 2 days post-stimulation cells were used for stimulation with MEL624 cancer cell line in the presence of soluble Siglec-5 Fc (10μg/ml) or control hIgG1 Fc (10μg/ml) protein. The assay was carried in the presence of Brefeldin A and Monensin. (**A**) Representative plots and summary of frequency of CD4 T cells expressing IL-2, IFN-*γ*, and TNF-*α* at 12hrs post-stimulation. (**B**) Representative plots and summary of frequency of CD4 T cells expressing CD107a. Statistical analysis: Ratio paired two-tailed t-test; * p<0.05. Each line represents an individual donor.

## DISCUSSION

In this study, we report that Siglec-5 is an inhibitory surface antigen for human T cell activation. Siglec-5 overexpression inhibits NFAT and AP-1 transcription factor activity. GBS virulence factor *β*-protein, a known Siglec-5 ligand, inhibits primary T cell cytokine production in a Siglec-5-dependent manner. Furthermore, a soluble form of Siglec-5 enhances cytokine production and granule release by cancer antigen-specific T cells. When taken together, our data demonstrate that Siglec-5 is a previously unrecognized inhibitory checkpoint molecule expressed by activated human T cells.

Our data show that surface expression of Siglec-5 by activated human T cells coincides with many other co-inhibitory and co-stimulatory molecules, including PD-1 and OX40 (Fig.1E and Fig.S3B). Other checkpoints, such as CTLA-4, 4-1BB, and ICOS are expressed by T cells earlier than Siglec-5, yet show similar overall expression kinetics (Fig.2F, Fig.S3A and S3C). Both co-stimulatory and co-inhibitory checkpoint molecules are known to play crucial roles in regulating T cell activation. In the case of co-stimulatory receptors, they synergize with TCR signaling and contribute toward the T cell effector functions. On the other hand, co-inhibitory receptors act to antagonize TCR signaling and prevent hyper-activation of T cells. These co-stimulatory and co-inhibitory opposing forces coming from the different co-stimulatory and co-inhibitory checkpoint receptors, whose activation depends on their ligand availability, define the fate of T cell activation, differentiation, and proliferation (*25*). Co-expression of multiple immune checkpoints and the difference in the timing of their ligand’s expression suggest that T cell responses can be differentially regulated based on the spatiotemporal availability of specific ligands expressed by antigen presenting cells. In lieu of this knowledge, we are currently investigating the identity and tissue distribution of molecules that bind Siglec-5 to determine potential ligand expression by normal and transformed cells.

It should be noted that the time course of Siglec-5 surface expression does not correspond to the level of total protein expression. In contrast to its surface expression at 48hrs post-stimulation (Fig.1C and 1D), we can detect Siglec-5 mRNA and total protein expression as early as 24hrs post-stimulation (Fig.2A and B). These data suggest that surface translocation and/or exo-/endocytosis of Siglec-5 is tightly controlled, similar to what has been observed for CTLA4 (17). Further studies are needed to elucidate how trafficking of Siglec-5 to the cell surface is regulated.

Mechanistically, the data we present in this work collectively suggest that Siglec-5 mediates its inhibitory function via the conserved ITIM and ITSM, in a manner that parallels PD-1 (*26, 27*). In support, previous studies on the Siglec-5 function in myeloid cells have shown that upon ligand engagement, ITIM and ITSM become phosphorylated, and that these phosphorylated motifs serve as recruitment sites for phosphatases such as Shp1 and Shp2 (*12, 13*) whose functional roles are to de-phosphorylate proteins involved in the membrane proximal signaling events. When we over-expressed Siglec-5 in Jurkat T cells, Siglec-5 significantly inhibited the activity of NFAT and AP-1 following TCR stimulation (Fig.3B). However, obtaining such a result in the absence of an added ligand to engage Siglec-5 is enigmatic and raised the question how Siglec-5 initiates its inhibitory effect. Intriguingly, our data also demonstrated that approximately 15% of Jurkat T cells express a putative ligand for Siglec-5 (Fig.S6B), suggesting that Siglec-5 was engaged either by the ligands expressed autologously or by neighboring Jurkat T cells. Unexpectedly, expression of a Siglec-5 mutant that lacks the ITIM and ITSM led to an increase of the NFAT activity in unstimulated cells. These data suggest that the ecto- and/or transmembrane domains of mutant Siglec-5 could interact with endogenous proteins, where presumably Siglec-5 inhibits their function in a dominant manner.

Our studies here have also shown that GBS β-protein, a specific ligand for Siglec-5, suppressed primary T cell activation in a Siglec-5 dependent manner. To accomplish this we used the N-terminal region of the β- protein, B6N (aa1-152), which is required and sufficient for engaging Siglec-5 (*28*), as a fusion protein with sfGFP (Fig.S4). Our data show that B6N::sfGFP fusion protein, but not control sfGFP, suppresses the production of proinflammatory cytokines such as IFN-γ and IL-22 by primary human CD4 T cells (Fig.4A). In contrast, B6N::sfGFP fusion protein caused a mild but reproducible increase in Th2 cytokines such as IL-4, IL-5, and IL-13, potentially due to the reduced IFN-γ production (29) (Fig.4B). Similarly, B6N::sfGFP fusion protein also reduced IFN-γ production by activated CD8 T cells (Fig.4A). Because IFN-γ plays a crucial role in the clearance of GBS in neonates (*30, 31*), our data here suggest that Siglec-5 mediated T cell suppression in neonates could be another mechanism by which β-protein expressing GBS modulates the immune response.

Our data showed that Siglec-5 interacts with a putative endogenous ligand(s) that inhibits the effector functions of human T cells and that multiple cancer cell lines express surface molecules that bind soluble Siglec-5 protein. Among the cell lines tested, the melanoma cell line MEL624 showed strongest binding to soluble Siglec-5 (Fig.S6). Given this observation, cancer cells could suppress the T cell response in part by engaging Siglec-5. Therefore, by blocking the ligand availability on cancer cells using soluble Siglec-5, we would expect a reinvigoration of T cell effector functions. Indeed, our data demonstrate that soluble Siglec-5 increased the frequency of tumor antigen specific CD4 T cells producing IL-2, IFN-γ, and TNF-α. In addition, we also observed an increase in the frequency of cells expressing CD107a (also known as LAMP-1) a lysosome-associated molecule that marks cells that have recently released cytotoxic granules (CGs). CGs are specialized lysosomes comprised of granzymes and perforins, which mediate targeted cell lysis when released extracellularly. When taken together, our data shown here suggest that soluble Siglec-5 can increase the ability of T cells to eliminate targeted cancer cells.

Immune checkpoint blockade therapies work by releasing the inherent inhibition of T cells to act on cancer, thereby allowing them to mount a robust immune response. To date, across fourteen different malignancies the FDA has approved seven different drugs targeting three different inhibitory checkpoints, namely PD-1, PD-L1, and CTLA-4. The approval of these therapies has revolutionized the way malignancies are treated and has brought hope for many patients. However, based on a 2018 study, even though approximately 43.6% of cancer patients in the US are predicted to respond to immune checkpoint therapies, the percentage of patients that actually mount a proper response is close to 12.5% (*32*). This low response rate is postulated to be due to either primary or acquired resistance by the tumors and suggests that blocking one inhibitory pathway is not enough to rescue the T cell response, as compensatory mechanisms by other checkpoint receptors could be upregulated to prevent T cell activation. As a result, extensive research has been focusing on identifying new checkpoint receptors. Currently, several immune checkpoint molecules are studied for potential use in therapies, including lymphocyte activation gene-3 (LAG-3) (*33*), T cell immunoglobulin and ITIM domain (TIGIT) (*34*), T cell immunoglobulin and mucin-domain containing-3 (TIM-3) (*35*), V-domain Ig suppressor of T cell activation (VISTA) (*36*), B and T cell lymphocyte attenuator (BTLA)(*37*) and B7 homolog 3 protein (B7-H3) (*38*). Our data show that Siglec-5 is another T cell checkpoint receptor that suppress the T cell responses. Future studies investigating the *in vivo* effect of Siglec-5 blockade will provide information about the potential use of Siglec-5 as a novel target of cancer immunotherapy.

A limitation of this study is the lack of *in vivo* data. The mouse ortholog of Siglec-5 is Siglec-F (*39*). However, besides sharing high sequence homology with Siglec-5, Siglec-F is closer to another CD33rSiglec, Siglec-8, with which it shares similar expression patterns, ligand preferences and functional outcomes (*40*). Thus, to study the T cell specific role of Siglec-5 in physiological or pathological conditions, a humanized mouse model needs to be established.

## MATERIALS AND METHODS

### Cell Culture

Complete RPMI used for suspension cell cultures was prepared with RPMI-1640 (HyClone, Logan, UT) and supplemented with 10% fetal calf serum (FCS) (Gemini Bioproducts, West Sacramento, CA), essential amino acids (Corning, Corning, NY), non-essential amino acids (Gibco, Waltham, MA), 1mM sodium pyruvate (Corning, Corning, NY), 50mM 2ME (Fisher Scientific, Waltham, MA), and 0.1M Hepes (Corning, Corning, NY) and Penicillin/Streptomycin (HyClone, Logan, UT). Adherent cells were maintained in DMEM supplemented with 10% FCS (Gemini Bioproducts, West Sacramento, CA) and Penicillin/Streptomycin (HyClone, Logan, UT). All cells were maintained with 5% CO_2_ at 37C.

Human lymphocytes were obtained from adult or cord blood. Adult peripheral blood mononuclear cells (PBMCs) were purchased from Key Biologics (Memphis, TN) or Zen Bio (Research Triangle Park, NC) and came from de-identified adult healthy donors. Cord blood mononuclear cells were isolated from whole umbilical cord blood kindly donated from Loyola University Medical Center from healthy donors (exclusion criteria: 1. autoimmunity; 2. malignancy; 3. use of immunosuppressive medication; 4. hyper or hypothyroidism). Heparinized blood was separated using Lymphopure density gradient medium (Biolegend, San Diego, CA) according to the manufacturer’s instructions.

Whole PBMCs were seeded at 1-2×10^6 cells per well in 48-well plates in the presence of 200ng/ml soluble anti-CD3 (OKT3 clone, Biolegend, San Diego, CA) and 10ng/ml IL-2 (Peprotech, Rocky Hill, NJ). For conditions extending past 3 days, cells were split 1:2 and fresh media supplemented with 10ng/ml IL-2 was added back.

Simian virus 40 large T antigen transfected Jurkat cells, a kind gift from Dr. Art Weiss (UCSF, San Francisco, CA), were maintained in complete RPMI, and were used for all transfection experiments.

MEL624 (ATCC, Manassas, VA) were maintained in DMEM (HyClone|Cytiva, Logan, UT) supplemented with 10% FCS (Gemini Bioproducts, West Sacramento, CA) and Penicillin/Streptomycin (HyClone, Logan, UT), and were used for antigen specific stimulation of T cells.

### T Cell Isolation and Culture

CD3+, naïve CD4+ and naïve CD8+ T cells were isolated from mononuclear cells from healthy adult or cord blood via negative selection using the MojoSort CD3+, naïve CD4+ or naive CD8+ T cell enrichment kit (Biolegend, San Diego, CA). Cell purity for all enrichments ranged between 90-98%. In all assays using enriched cells, cells were cultured at 0.5-1×10^6 cells/ml in plates coated with anti-CD3 (OKT3, 5ug/ml) and anti-CD28 (5ug/ml) and complete RPMI media supplemented with 10ng/ml IL-2.

### Flow Cytometry

Antibodies used for the flow cytometry analysis were anti-CD4, -CD8, -Siglec 5 (1A5), - Siglec-3, -Siglec-7, -Siglec-9, -Siglec-10, -CD137, -PD-1, , -Granzyme B, -OX40, -ICOS, - CTLA-4, -IFN-*γ*, -TNF-*α*, -IL-2, -CD107a, (Biolegend, San Diego, CA). Cells were blocked with human Fc receptor blocking solution (Human TruStain FcX, Biolegend) for 10 minutes on ice. For detection of cell surface markers, cells were stained with antibody cocktail for 30 minutes at 4C. Intracellular markers were analyzed after fixing and perming with 4% paraformaldehyde and permeabilization buffer (50mM NaCl, 0.02% NaN3, 5mM EDTA, 0.5% TritonX, pH7.5), respectively. Data was collected using BD FACSCanto or BD LSRFortessa and analyzed using FlowJo v.10 software.

### Western Blot and Immunoprecipitation Analysis

For Western blot analysis, cells were lysed in non-reducing SDS sample buffer (2% SDS, 0.05% bromophenol blue, 62.5mM Tris-HCl, pH 6.8, 10% glycerol, 10mM Iodoacetamide (IAM)) and boiled 2 x 5min. Equal amount of proteins, based on cell number, was loaded, and separated on SDS-PAGE gel. PVDF membrane transferred proteins were probed with antibodies against anti-Siglec 5/14 (clone 1A5, Biolegend, San Diego, CA).

For immunoprecipitation of Siglec-5, cord blood mononuclear cells were stimulated using soluble anti-CD3 (200ng/ml) and IL-2 (10ng/ml) for 2-3 days. 50-60×10^6 cells were lysed in 1ml of lysing buffer containing 0.5% NP-40, 0.15M NaCl, 5mM EDTA, and protease and phosphatase inhibitors. Cell lysates were pre-cleared with Protein G-Sepharose pre-incubated with mIgG1 isotype antibody for 2hrs, and immunoprecipitated with CNBr-activated Sepharose conjugated with anti-Siglec5/14 antibody (clone 1A5) for 2hrs. After washing 3 times with lysis buffer, the immunoprecipitants were either boiled in 1x SDS buffer (2% SDS, 0.05% bromophenol blue, 62.5mM Tris-HCl, pH 6.8, 10% glycerol, 125mM DTT) or further subjected to de-glycosylation with PNGase-F (NEB, Ipswich, MA) following the manufacturer’s instructions. De-glycosilated samples were also boiled in reducing SDS buffer. Samples were run on SDS-PAGE gel, and proteins were transferred on PVDF membrane. Membranes were probed with antibodies against anti-Siglec5/14 (polyclonal antibody, R&D, Minneapolis, MN).

### Reporter and Expression Constructs

cDNA sequences for Siglec-5 (SinoBiological) and Siglec-14 (IDT, inc.) were subcloned into a pME vector using PCR based cloning. To generate a truncated version of Siglec-5 (pME- tSiglec-5) the ITIM and ITSM were excised from the pME-Siglec 5 plasmid with the restriction enzymes BbsI/MfeI (NEB, Ipswich, MA) and blunt ends were generated with DNA Polymerase I, Large (Klenow) Fragment (Thermo Fisher, Walthman, MA). The blunt ended plasmid was then ligated using T4 DNA polymerase (Thermo Fisher, Walthman, MA). NFAT-luciferase and AP-1 luciferase were previously described(*41*). All plasmid DNA was prepared using CsCl purification method.

### Luciferase Assay

Jurkat Tag cells (2×10^6) were transfected using electroporation (0.8uF and 0.260mV) with 10ug of NFAT or AP-1 reporter DNA, along with 1ug of cytomegalovirus promoter-driven expression vector for *Renilla* luciferase and 30ug of the expression vectors for pME-Siglec-5 or pME-*Δ*Siglec 5. 48hrs post transfection equal number of cells were plated on anti-CD3 coated wells in 96-well plates. 4hrs post stimulation cells were harvested, and luciferase activity was measured using a bioluminescent reporter assay kit (Promega, Madison, WI) and a luminometer. For each transfection condition, the relative luciferase activity was adjusted based on Renilla luciferase activity.

### Cloning, expression and purification of B6N::sfGFP and sfGFP

The DNA sequence encoding the B6N region (amino acids 1-152) of the Group B Streptococcus *β*-protein(*12*) was codon-optimized for expression in *E. coli* and synthesized as a gene fragment (Integrated DNA technologies, Coralville, IA). The B6N region was then fused to a super-folder green fluorescent protein variant, sfGFP(*42*), with a flexible linker (G-S-G-G-G-G-S-G-G-G-G-S) by overlap-extension PCR. The resulting B6N-linker-sfGFP sequence, or the control linker-sfGFP sequence, was amplified for use in a restriction-less cloning method as previously described(*43*). The resulting DNA fragments were ligated into pET15DG1(*44*) digested with NdeI and BamHI.

6xHis-tagged proteins were expressed in T7 Express *lysY/I^q^ E. coli* cells (New England Biolabs, Ipswich, MA). Cultures were grown at 37 °C in LB medium supplemented with 50 μg/mL carbenicillin. Protein expression was induced at an OD_600_ of 0.8 with 1 mM IPTG. The temperature was then reduced to 18 °C and the cultures were grown overnight with shaking at 150 rpm. Cells were then harvested by centrifugation, resuspended in 25 mL buffer A (50 mM Tris-CL pH 8.0, 150 mM NaCl, 25 mM imidazole, 1 mM BME) per liter of culture. Following supplementation with 1 mM PMSF, cells were lysed by sonication using a Branson Sonifier S-450 cell disruptor (Branson Ultrasonics Crop., Danbury, CT) for a total of 5 minutes of sonication at 50% amplitude with 30 second pulses, separated by 1 minute rest periods. The lysate was then clarified by centrifugation at 16,000 x g for 45 minutes and then applied to 3 mL Ni-NTA resin (Qiagen, Valencia, CA) equilibrated with buffer A. After washing the column with buffer A, 6xHis-tagged protein was eluted in buffer B (50 mM Tris-Cl pH 8.0, 150 mM NaCl, 250 mM imidazole, 1 mM BME). Protein-containing fractions were analyzed by SDS-PAGE, pooled, and dialyzed against buffer C (50 mM Tris-Cl pH 8.0, 150 mM NaCl, 1 mM BME). Proteins were further purified by Size Exclusion Chromatography (SEC). SEC experiments were carried out using an AKTA Pure FPLC system (General Electric, Boston, MA) equipped with a HiLoad 16/600 Superdex 75 pg column equilibrated in buffer C. Samples (2 mL) were injected onto the column and separated at a flow rate of 0.5 mL/min. 1 mL fractions were collected and analyzed by SDS-PAGE. The desired fractions were pooled and dialyzed into storage buffer (50 mM Tris-Cl pH 8.0, 150 mM NaCl, 40% glycerol, 1 mM DTT). Endotoxin was removed using Pierce High Capacity Endotoxin Removal Resin (Fisher Scientific, Waltham, MA). Endotoxin was quantified using a Pierce LAL chromogenic Endotoxin quantitation kit (Fisher Scientific, Waltham, MA). Proteins were stored at -80 °C.

### B6N::sfGFP ELISA

Purified B6N::sfGFP or sfGFP were immobilized on Nunc Maxisorp 96-well plate (Thermo Fisher Scientific, Waltham, MA) for 2 hours at RT. Target coated plates were washed 3x with PBST (0.05% Tween-20) and then blocked with 3% BSA for 1 hour at RT. After washing 3x with PBST, recombinant Siglec 5-Fc chimeric protein (Biolegend, San Diego, CA) was added and incubated for 2hrs at RT, followed by 3x washes and incubation with anti-human IgG-HRP antibody for 1hr. After 3xPBST washes, 1-Step Ultra TMB-ELISA (Thermo Fisher Sciebtific, Waltham, MA) was added and absorbance was determined at 450nm with a microtiter plate spectrophotometer (BioTech).

### T Cell Stimulation with B6N::sfGFP

Naïve CD4+ or CD8+ T cells were cultured with plate bound anti-CD3 (5ug/ml) and anti-CD28 (5ug/ml) stimulation in the presence of IL-2 (10ng/ml) for 3 days. Cells were than harvested, washed and re-stimulated with plate-bound anti-CD3 (5ug/ml), anti-CD28 (5ug/ml) and B6N::sfGFP or sfGFP (166nM) for additional 3 days. To reverse the inhibitory effects B6N::sfGFP has during T cell stimulation, equimolar amount of B6N::sfGFP or sfGFP were pre-incubated with recombinant hIgG1 Fc or Siglec-5-Fc proteins for 1hr at 4C with rotation. The proteins were than coated on 96 well non-tissue cultured plates along with anti-CD3 and anti-CD28, before adding the 3 days stimulated CD4 T cells. Supernatants from 3-day long re-stimulated cultures were collected and used for cytokine bead array (CBA) analysis for the expression of IL-5, IL-13, IL-2, IL-6, IL-9, IL-10, IFN-*γ*, TNF-*α*, IL-17A, IL-17F, and IL-22 using the LEGENDplex Human Th Cytokine (Biolegend, San Diego, CA). Cells from 3-day long re-stimulated cultures were used for evaluating expression of: CD4, CD8, Siglec-5 and Granzyme B.

### Evaluation of Putative Siglec-5 Ligands Expressed by Cell Lines

Cell lines: MEL624, Jurkat, Raji, THP1, U937, T2, SNU-475, HEK293t, A549 were blocked with human Fc receptor blocking solution (Human TruStain FcX, Biolegend) for 10 minutes on ice. Cells were than stained with Siglec-5 Fc chimeric protein (Biolegend, San Diego, CA) pre-incubated with anti-human IgG Fc fluorescent antibody (Biolegend, San Diego, CA). As a negative control, cells were stained with anti-human IgG Fc antibody alone.

### Human 1383i TCR+ T Cell Stimulation

Human 1383i TCR+ T cells are tyrosinase specific and HLA-A2 restricted T cells that are generated by retroviral transduction of activated primary human T cells with the 1383i T cell receptor (isolated from tumor infiltrating lymphocytes from a melanoma patient) (*45*). To stimulate the 1383i TCR+ T cells, T2 (1×10^6 cells/ml) (ATCC, Manassas, VA) cells were pulsed with tyrosinase peptide 368-376 (YMDGTMSQV) (10ug/ml) for 2hrs at 37C. T2 pulsed with tyrosinase were than irradiated with 4425cGy and used for culturing with 1383i TCR+ T cells at a ratio of 3:1 (1383i T cells : T2). After 2 days of stimulation, 1383i TCR+ T cells were harvested, and cell activation and expression of Siglec-5 were verified. Cells were than used for re-stimulation with MEL624 (ATCC, Manassas, VA) at 1:1 ratio. Before co-culture, MEL624 was pre-treated with hIgG1-Fc (10ug/ml) (Biolegend, San Diego, CA) or Siglec-5-Fc chimeric protein (10ug/ml) for 30mins on ice. Cell cultures were carried in the presence of 1x Monensin and 1x BrefeldinA (Biolegend, San Diego, CA). At 12hrs post-stimulation, expression of CD107a, IFN-*γ*, TNF-*α* and IL-2 was evaluated in the CD4 and CD8 compartments of 1383i TCR+ T cells.

## Supplementary Materials

**Fig. S1.**
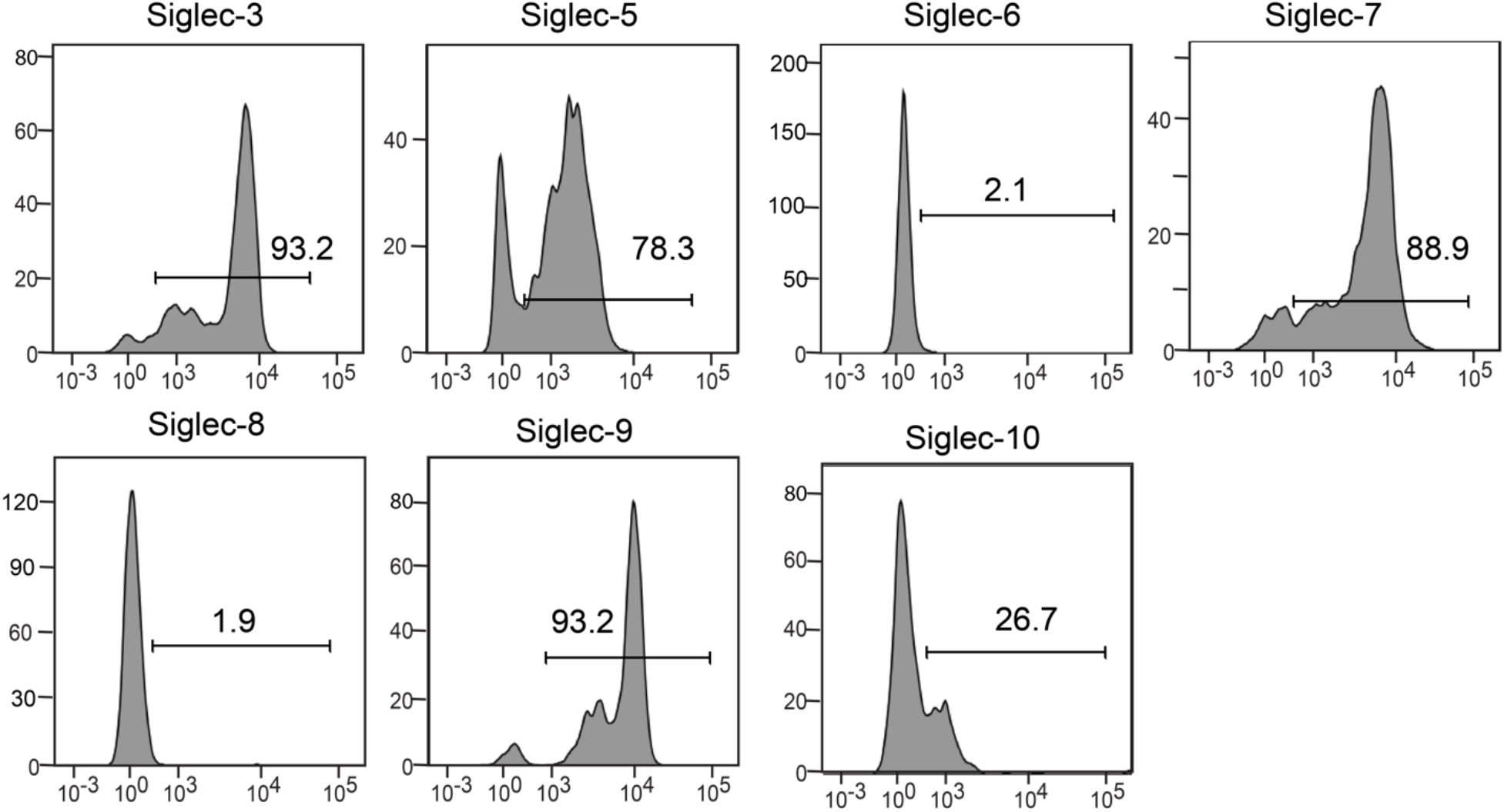
Monocyte expression of CD33rSiglecs. Freshly isolated PBMCs were stained for Siglec-3, -5, -6, -7, -8, -9 and -10. Expression levels for each Siglec by the monocytes fraction of cells were evaluated by flow cytomtery. Ab and IL-2. On day 2 post stimulation, co-expression of Siglec-5 and (**A**) PD-1 and (**B**) CTLA-4 were evaluated by flow cytometry.

**Fig. S3.**
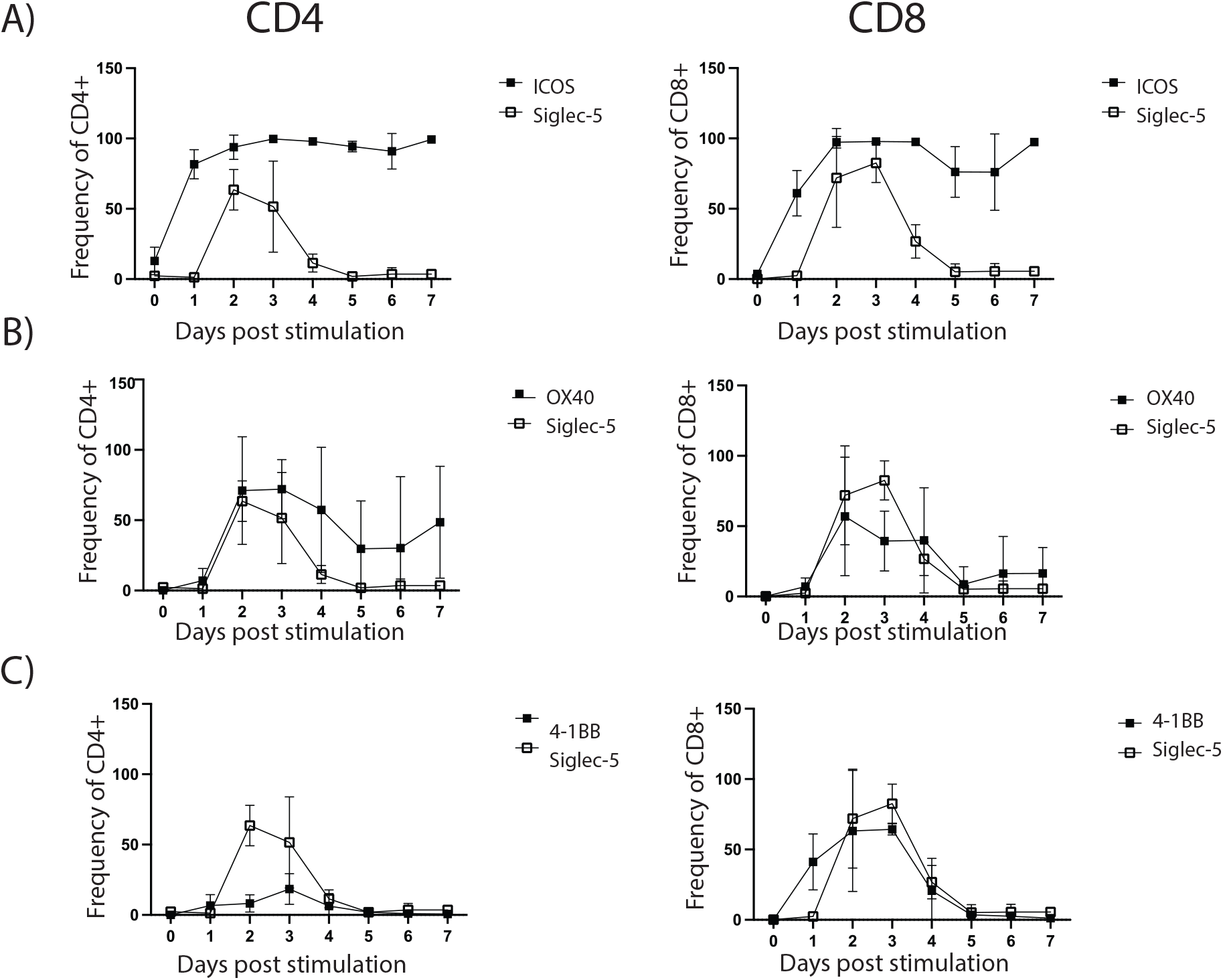
Expression kinetics of Siglec-5 and co-stimulatory checkpoint receptors in T cells. Cord blood mononuclear cells were stimulated with soluble anti-CD3 and IL-2 for up to 7 days. Cells were split every 2-3 days. At each time point, cells were stained and analyzed for expression of Siglec-5 and (**A**) OX-40, (**B**) ICOS and (**C**) 4-1BB in CD4 and CD8 T cells.

**Fig. S4.**
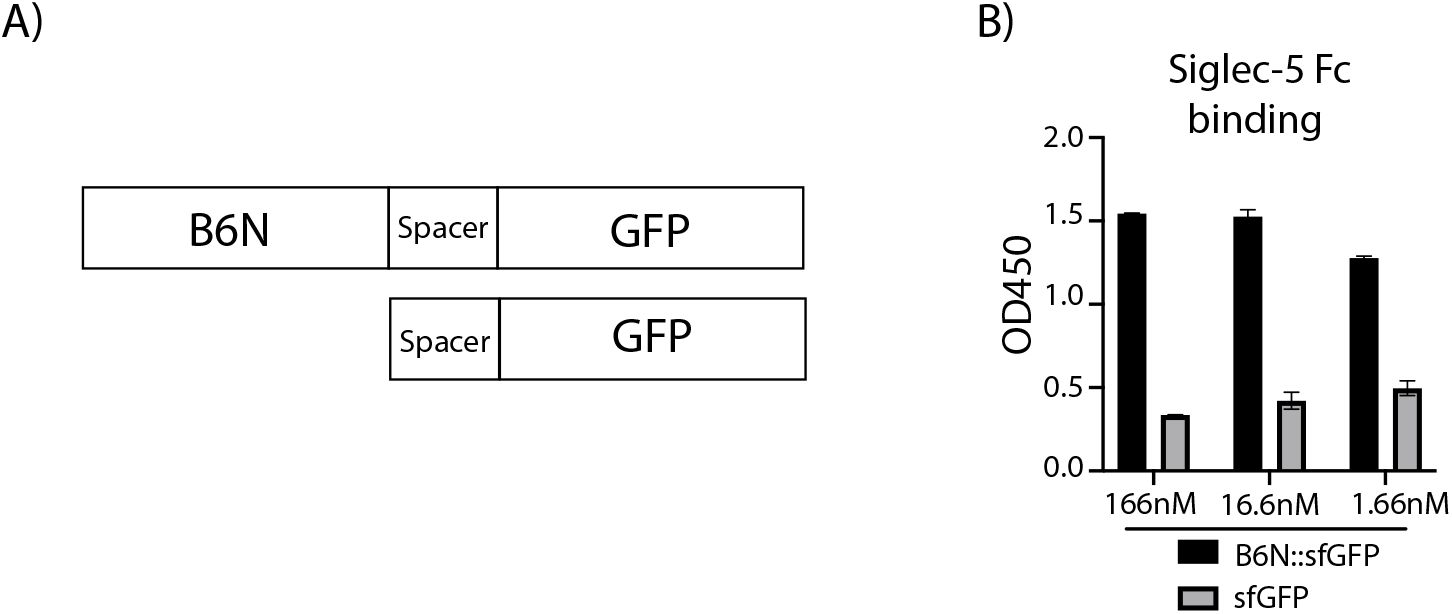
Siglec-5 binding to B6N::sfGFP. (**A**) Schematic representation of B6N::sfGFP and sfGFP proteins. (**B**) Graded concentrations of B6N::sfGFP (black bars) and sfGFP (gray bars) were immobilized on ELISA plates. Unconjugated recombinant Siglec-5 Fc chimeric protein was used as detection reagent. HRP-conjugated anti-hIgG was used for detection of Siglec-5 Fc binding.

**Fig. S5.**
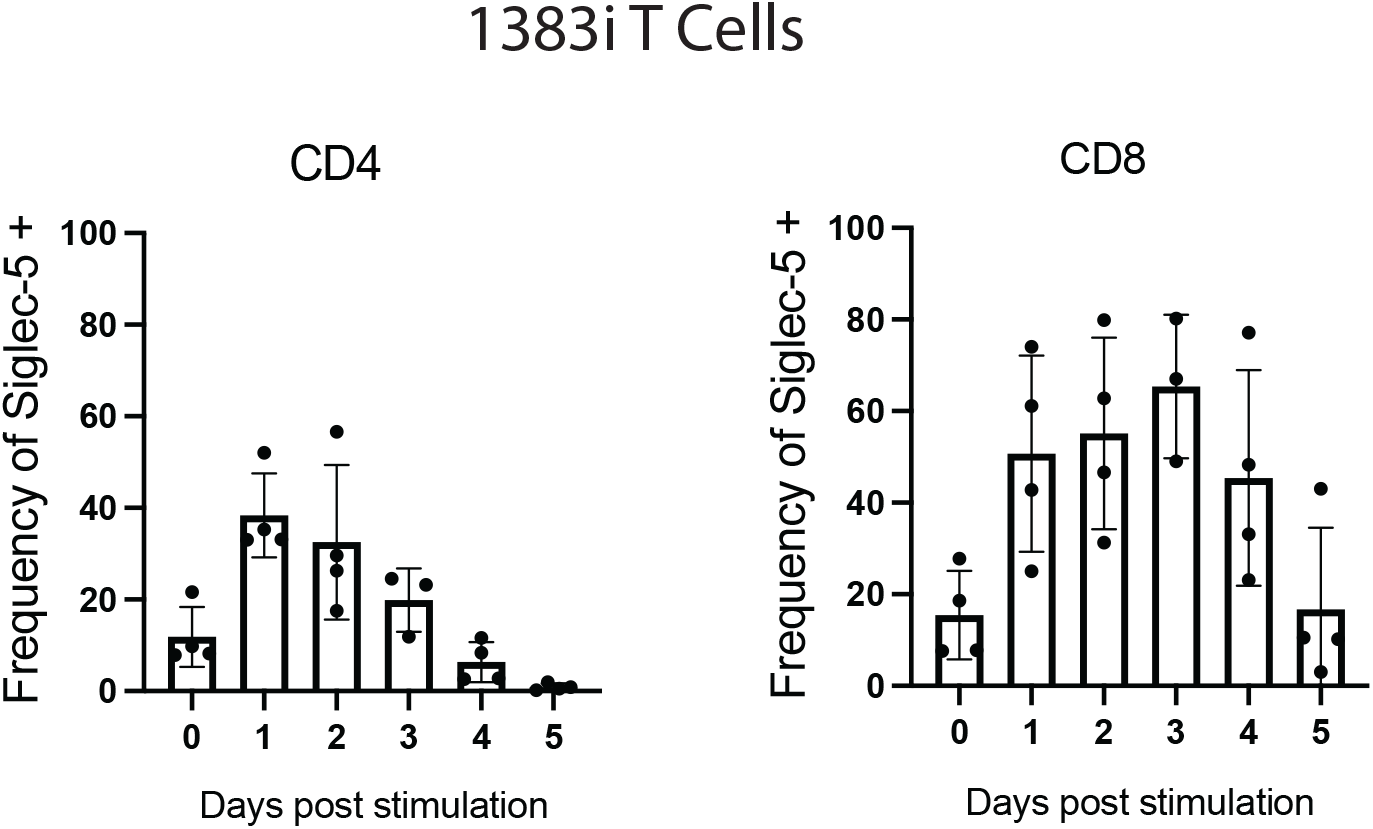
Siglec-5 expression on tyrosinase specific T cells. Adult T cells were transduced with 1383i TCR specific for the melanoma antigen tyrosinase. Transduced T cells were enriched using the CD34t selection marker which is expressed in-frame with the tyrosinase specific 1383i TCR and denotes 1:1 expression ratio. 1383i TCR+ T cells were than stimulated with a 1:3 ratio of T cells to irradiated T2 pulsed with tyrosinase, and cultured for 0, 1, 2, 3, 4, and 5 days. Siglec-5 expression within the CD4 and CD8 compartment was evaluated at each time point.

**Fig. S6.**
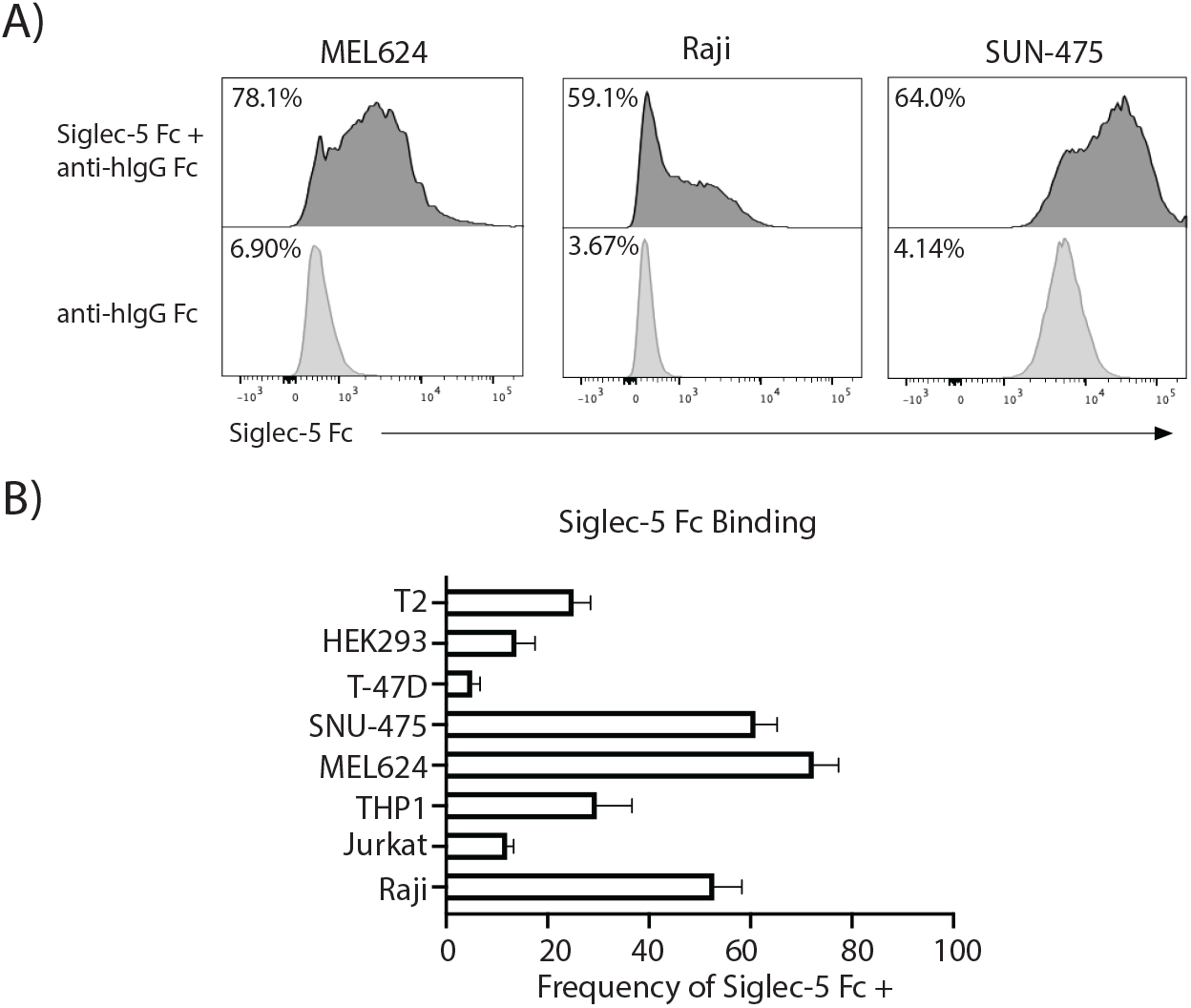
Siglec-5 Fc binding to cancer cell lines. (**A**) Representative plots of human cancer cell lines stained with recombinant Siglec-5 Fc chimeric protein and anti-human IgG1 Fc fluorescent antibody. (**B**) Summary of multiple staining for different cell lines by Siglec-5 Fc protein. Percentage of cells that stained positive are shown.

## Acknowledgments

We thank Pat Simms at the FACS Core Facility (Loyola University Chicago) for assistance with FACS. We thank Yi Wei Lim and Christina Cunha (Loyola University Chicago) for helpful discussions and insightful criticism of the data. We thank the stuff and patients at the Labor and delivery unit at Loyola University Medical Center for the kind donation of cord blood.

## Author contributions

Conceptualization: AV, MI

Methodology: AV, DG, GS, PW

Investigation: AV, DG

Visualization: AV

Funding acquisition: MI, AU, AV

Project administration: MI

Supervision: MI, AU, MN, PW

Writing – original draft: AV

Writing – review & editing: AV, MI, AU

## Data and materials availability

All data are available in the main text or the supplementary materials.

